# Neural correlates of category learning in monkey inferior temporal cortex

**DOI:** 10.1101/2023.12.05.568765

**Authors:** Jonah E Pearl, Narihisa Matsumoto, Kazuko Hayashi, Keiji Matsuda, Kenichiro Miura, Yuji Nagai, Naohisa Miyakawa, Takafumi Minanimoto, Richard C Saunders, Yasuko Sugase-Miyamoto, Barry J Richmond, Mark A G Eldridge

## Abstract

We trained two monkeys implanted with multi-electrode arrays to categorize natural images of cats and dogs, in order to observe changes in neural activity related to category learning. We recorded neural activity from area TE, which is required for normal learning of visual categories based on perceptual similarity. Neural activity during a passive viewing task was compared pre- and post-training. After the category training, the accuracy of abstract category decoding improved. Specifically, the proportion of single units with category selectivity increased, and units sustained their category-specific responses for longer. Visual category learning thus appears to enhance category separability in area TE by driving changes in the stimulus selectivity of individual neurons and by recruiting more units to the active network.

## Main

Area TE (TE) is the most rostral brain area of the ventral visual stream^1^. Lesion studies have demonstrated that TE is essential for the accurate categorization of perceptually ambiguous stimuli^2,3^. Previous studies of TE’s role in visual categorization have demonstrated that, after extensive behavioral training, neurons in TE become tuned to the diagnostic features of parameterized stimuli^4–6^. However, monkeys – like humans – can rapidly learn new category groupings, and subsequently generalize those categories to new exemplars, without the need for extensive training^7^. Here, we examined neural correlates of rapid visual categorization of unparameterized natural images, while simultaneously recording from large numbers of single units in TE. We show that the recruitment of additional units to the network results in more accurate coding that correlates with improved behavioral performance.

Two Japanese monkeys (monkeys ‘R’ and ‘X’) were trained to categorize natural images of cats and dogs (Figure 1; see Methods and Supp Fig 1 for task design). Monkey X took one session and Monkey R took three sessions to learn to categorize 40 training images presented one at a time (10-50 trials per image per session). The monkeys were then tested on a larger, trial-unique set of cats and dogs (Supp Fig 2A), which they categorized with above-chance accuracy (∼70%) (although this was markedly below their peak accuracy of ∼90% during the training phase). The accurate responding to trial-unique stimuli is evidence that the monkeys used a generalization strategy (i.e., categorization) to discriminate the trial-unique set, as opposed to a memorization strategy. Images were cropped onto white backgrounds and had similar low-level visual statistics such as hue and saturation (Supp Fig 2), so it is unlikely that the monkeys distinguished the cats and dogs by any method other than integrating along the multiple visual dimensions that, combined, allow for reliable discrimination between the two species. We selected cats and dogs as the testing categories because we have previously demonstrated that these are more perceptually challenging to discriminate than alternative pairs of categories, such as cars vs. trucks, or human faces vs. monkey faces^2^.

**Figure 1:**
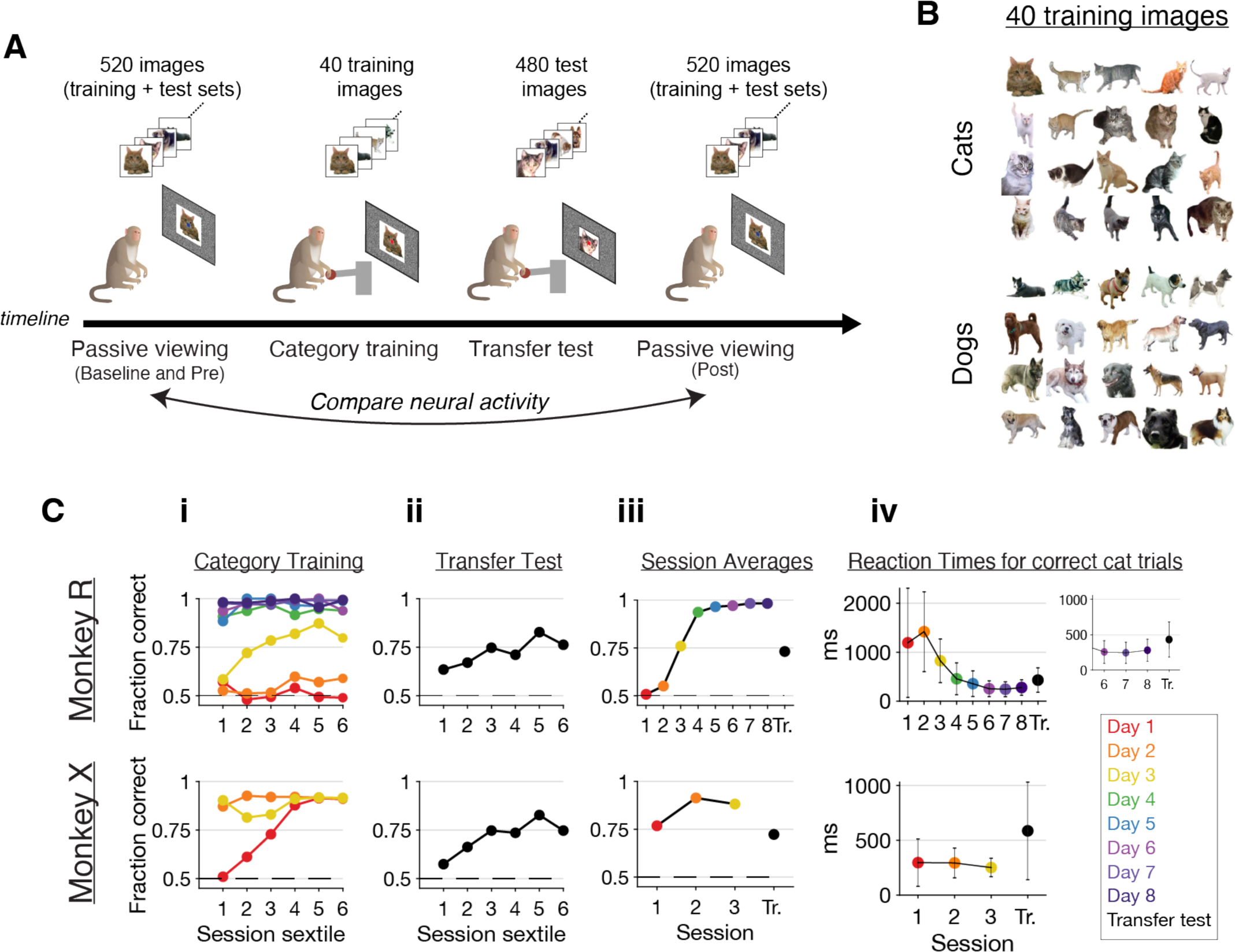
**A,** schematic of the experimental timeline. Neural activity was compared before and after monkeys learned to categorize natural images of cats and dogs. Monkeys were trained with 40 images and tested using 480 similar held-out images. All passive viewing sessions used all 520 images, randomly interleaved in blocks. **B,** the 40 images used for training. **C,** behavioral data from the category training, colored by session number; **i**, fraction correct trials split by session sextile; **ii**, same as **i** but for the first 480 completed trials of the transfer testing session, in which monkeys had only one opportunity to categorize each test image; **iii**, session averages for the data shown in **i** and **ii**; **iv**, reaction times for correct cat trials (release-on-red trials, see Methods), which correspond approximately to the time it takes monkeys to categorize the image^3^. Note the increased reaction times on the transfer test day, indicating non-expertise.

To investigate whether and how category learning changed the stimulus-evoked activity of neurons in TE, we recorded neural activity in TE while the monkeys passively viewed images of cats and dogs either before or after the category training. In total, we recorded 348 single units before training (148 from Monkey R and 200 from Monkey X) and 333 after training (139 and 194, respectively), from three chronically implanted Utah electrode arrays implanted in each subject (96 electrodes per array; Fig 2A). On each trial, monkeys fixated on a central point and five cat and/or dog images were presented sequentially (350-400 ms stimulus duration and inter-stimulus interval). During these passive viewing sessions, monkeys were rewarded following the presentation of all five images if they maintained fixation throughout the trial. Passive viewing recordings were performed at three times relative to the category training: for a week in the period a few months before the dog-cat discrimination training (“baseline”); for one day directly before the training (“pre-training”); and for one day directly after (“post-training”).

**Figure 2:**
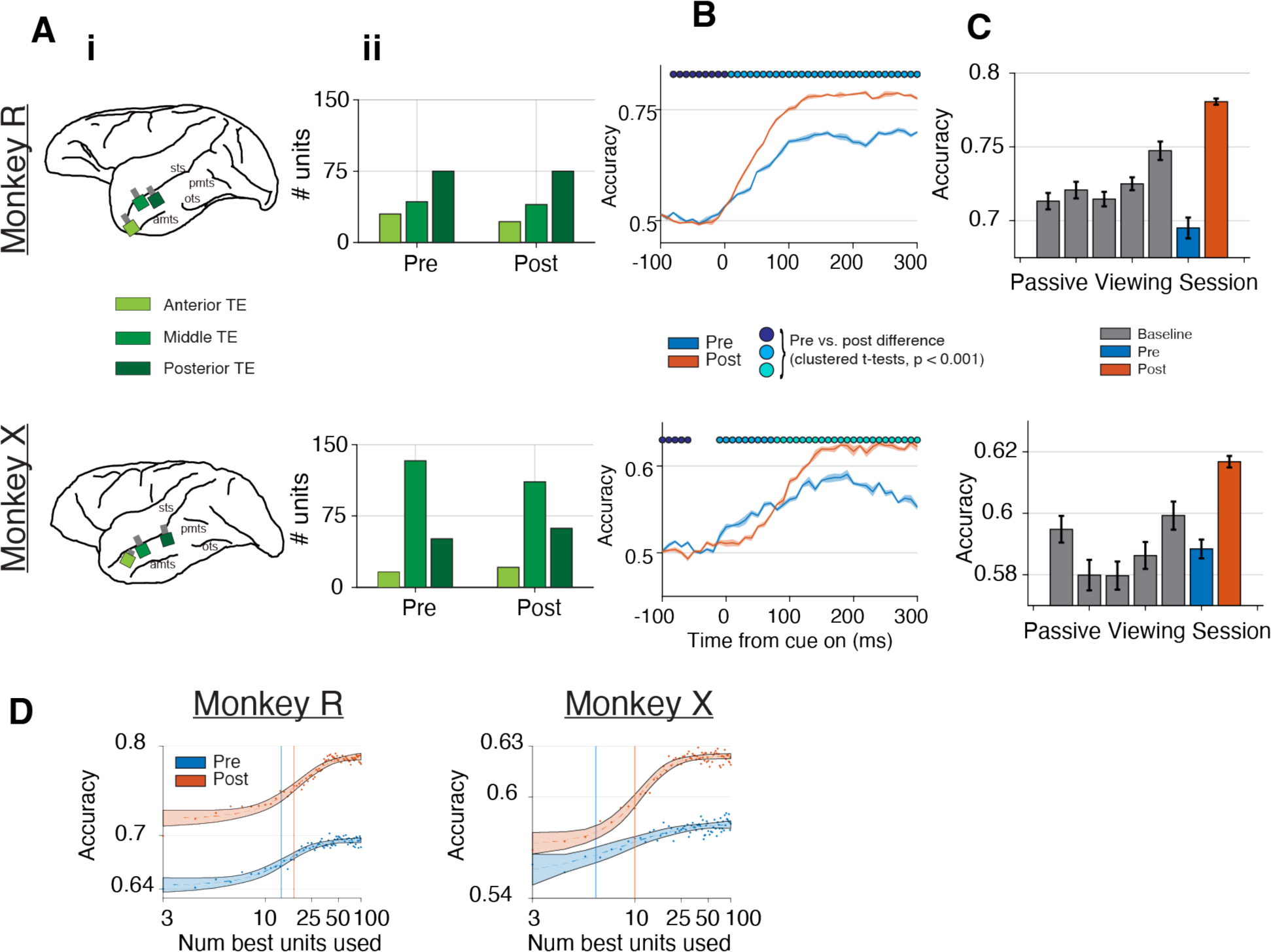
**A, i** Utah array locations in both monkeys, **ii** number of single units recorded from each pre/post-training session, from each Utah array. **B,** time-course of abstract category SVM decoders (see Methods), trained on neural population response vectors (spike counts in each 100 ms bin) (mean +/- shaded s.e.m.). Bouts of significant pre- vs. post-training difference were determined with t-tests and a cluster-based permutation procedure that uses trial-shuffled spike counts^9^. **C,** accuracy of the abstract category SVM decoders in the 175-275 ms bin, across experimental sessions (mean +/- s.e.m.). **D,** accuracy of abstract category SVM decoders in the 175-275 ms bin, with increasing numbers of the top 100 units used for training (see Methods). Broken vertical lines: half-maximal accuracy for pre- (blue) and post-training (red), respectively. See Supp Fig 3C for sigmoid parameters.

When comparing neural activity from the pre- and post-training days, linear support vector machines (SVMs) trained to decode image category from population vectors of spike counts (“abstract category SVMs,” 100 ms window size, see Methods) more accurately predicted category on the post-training day, from both the full neural populations combined across all three electrode arrays (Fig 2B) and all individual array subpopulations (Supp Fig 3B) (p < 0.05, cluster-based permutation corrected for multiple comparisons; see Methods). The “abstract category” decoding method holds out a set of images from training, forcing the decoder to rely on category-level information to subsequently predict the category of those images, rather than learning the category of each individual image. Decoding accuracy displayed the same trends for all bin sizes, and plateaued 175-275 ms after stimulus onset (Supp Fig 3A). Decoding accuracy at this plateau (spike count window 175 – 275 ms after stimulus onset) increased between pre- and post-training days by 8.6 and 2.8 percentage points for Monkey R and Monkey X, respectively (Fig 2C). The magnitude of this effect is comparable with previous studies of visual learning in TE^5^. Similar effects were also seen when the SVM input was restricted to only the 480 trial-unique images, which the monkeys had only 1-5 opportunities to see on the transfer test day (as opposed to 100+ opportunities across multiple days with the 40 training images) (data not shown). It seems unlikely that the monkeys would have memorized 480 initially trial-unique images and their reward associations in one exposure to the set. Thus, this category training enhanced visual category coding in area TE.

To determine what fraction of single units contributed to the observed population-level category coding, we re-trained the decoders on different-sized subpopulations, adding units in order of their decoding accuracy in a one-dimensional decoder (Fig 2D). As single units were added, decoding accuracy increased to an asymptote. Sigmoidal fits showed that the number of units needed for half-maximal accuracy increased post-training (13.3 to 17.0 units, p <0.05; and 5.8 to 9.8 units, p > 0.05), as did the amplitudes and midpoint-slopes of the fits (Supp Fig 3C). These results suggest that the population-level changes involved recruiting more units to the category-processing network, and were not simply due to a few single-units becoming strongly category selective.

Using a GLM to model the effect of category on the neural responses for each neuron confirmed that a larger proportion of single units responded selectively to category post-training when compared to pre-training (175-275 ms window; Monkey R, 34% to 55% of neurons, p = 2.3e-4, chisq = 13.6, df=1; Monkey X, 32% to 45% of neurons, p=8.8e-3, chisq = 6.9, df=1) (Figure 3A). This increase was observed in all three array locations, though not all increases reached significance (Supp Fig 4A). As in the decoding analysis, the changes observed in the GLM results between the first and last baseline days were smaller than those seen across training (except in the anterior array of Monkey R; Supp Fig 4B), arguing against a role for familiarity effects in driving the observed results.

**Figure 3:**
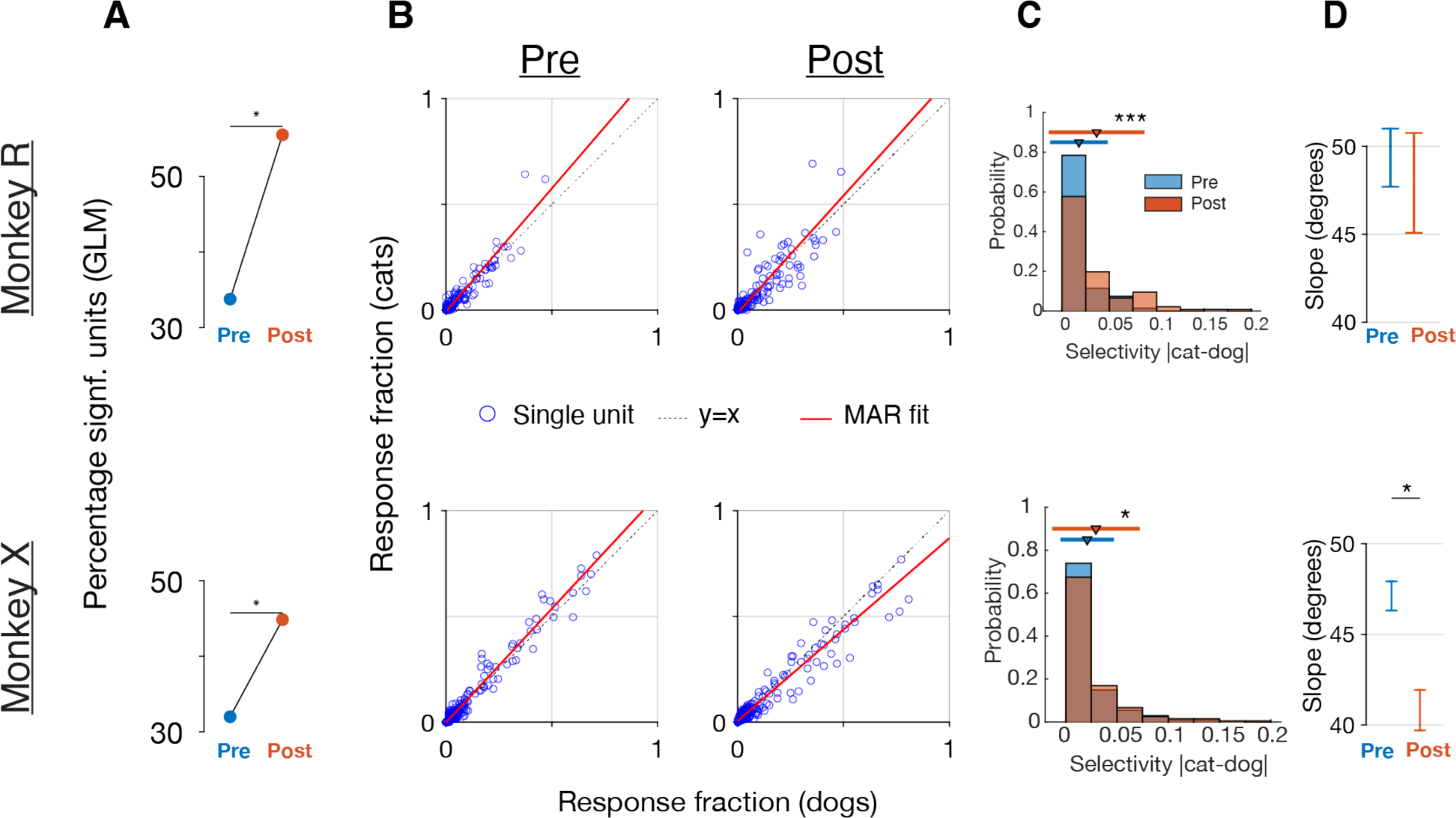
**A,** fraction of single units significant in a GLM regressing image category vs. spike count 175-275 ms after image onset. **B,** all single units’ responsiveness to cats or dogs, across sessions, as measured by the fraction of images from each category that evoked a significant visual response. Black dotted line, unity; red line, best-fit line from major-axis regression. Slopes of fit lines = 1.20 and 1.06, Pearson’s correlations = 0.95 and 0.98. **C,** distributions of the category selectivity shown in B, summarized for each unit by the absolute difference between the fraction of cat and dog images evoking a significant response. Triangle and line above the histograms represent mean and std, respectively. *, p < 0.05; ***, p < 0.001. **D,** estimated slope (from major-axis regression; mean + 95% confidence intervals) for best-fit lines in **b**.

To determine to what extent changes in single-unit image selectivity drove these effects, we calculated the proportion of cat and dog images to which each unit was significantly visually responsive (spike counts from 175 to 275 ms after stimulus onset vs. -150 to -50 before stimulus onset, paired t-tests, p < 0.05). Image selectivity before training was sparse, with 21% and 23% of units responding to more than 10% of images, but only 5% and 12% of single units responding to more than 25% of the images, for Monkeys R and X, respectively. This proportion increased moderately after training, with 34% and 28% responding to more than 10% of images (p=0.01 chisq=6.4, df=1; and p=0.27 chisq=1.2, df=1, respectively); and 10% and 18% of units responding to more than 25% of images (p=0.13, chisq=2.3, df=1; and p=0.12, chisq=2.4, df=1, respectively). These increases, as well as the total post-training proportions, were larger than any observed during baseline testing (21% +/- 3% and 13% +/- 1% units [mean +/- std] responding to >10% of images on baseline days, respectively). Single units were almost equally responsive to cats and dogs, with a mean absolute between-category difference of ∼2%. This difference, a rough proxy for single-unit category selectivity, increased slightly but significantly in both monkeys after training (p < 0.05, unpaired t-test; Figure 3B,C), reaching higher mean selectivity than any observed during baseline testing (3.7% mean difference post-training vs. 1.8% pre-training and 2.4% +/- 0.3% over baseline days; and 2.9% mean difference post-training vs. 2.1% pre-training and 1.4% +/- 0.3% over baseline days, respectively). Additionally, we observed a small but consistent decrease in single-unit sparseness^8^ after category training (p <= 0.05 for sparseness; p < 0.05 and p=0.08 for Nst; one-sided ranksum tests) (Supp Fig 4D); that is, the neurons responded to a larger number of images, and with more evenly distributed spike counts, after training. A greater number of neurons shifted their responsiveness toward dogs — the rewarded category— in both monkeys, although the shift toward dogs only reached statistical significance in Monkey X (no overlap/overlap, respectively, of analytical 95% confidence intervals for slope of the major-axis regression; Figure 3D). This shift in responsiveness for Monkey X was far greater than any changes observed during baseline testing (data not shown). We noted a similar effect in the coefficients from the GLM model, in which more units tended to have dog-preferring than cat-preferring coefficients after training, though only the coefficients from Monkey X reached significance (p = 0.1 and p < 0.001, respectively, two-sided ranksum test) (Supp Fig 4C). We did not observe a difference between the sparseness of responses to cats versus that to dogs either before or after training (p > 0.05, ranksum test), nor were the absolute spike rates for the significantly category-coding units higher post-training (p > 0.05, ranksum test), arguing against a non-specific spike rate increase due to reward association. We conclude that category training broadened single-unit image responses to one category or the other; that is, some units became more broadly tuned to dogs, while others became more broadly tuned to cats.

To assess whether the population encoding of category identity is stable or dynamic within a given trial, we applied a decoding analysis in which we trained a classifier using data from one time period and tested the classifier using data from a different time period (Figure 4A). Category information could be decoded from the population almost equally well at all time points from 75 ms to 450 ms after stimulus onset, regardless of the data used for training within the same monkey, indicating a relatively static (stable) representation of category identity over this timeframe. To better understand how this encoding is supported at the single unit level, we visualized the time course over which significant differences to category emerged for each unit (see Methods) (Figure 4B). Category selectivity in single neuron responses developed over the course of ∼400 ms from stimulus onset, with the majority of neurons beginning to show selectivity from 100 – 200 ms (see Supp Fig. 5 for smoothed PSTHs of example neurons). After training, units displayed longer periods of continuous category coding (Fig 4C). Additionally, the proportion of units with significant category selectivity increased on four of the six arrays (the anterior array of Monkey X also showed an increase but did not reach statistical significance; the amount of data was small, causing the test to be underpowered) (Figure 4D).

**Figure 4:**
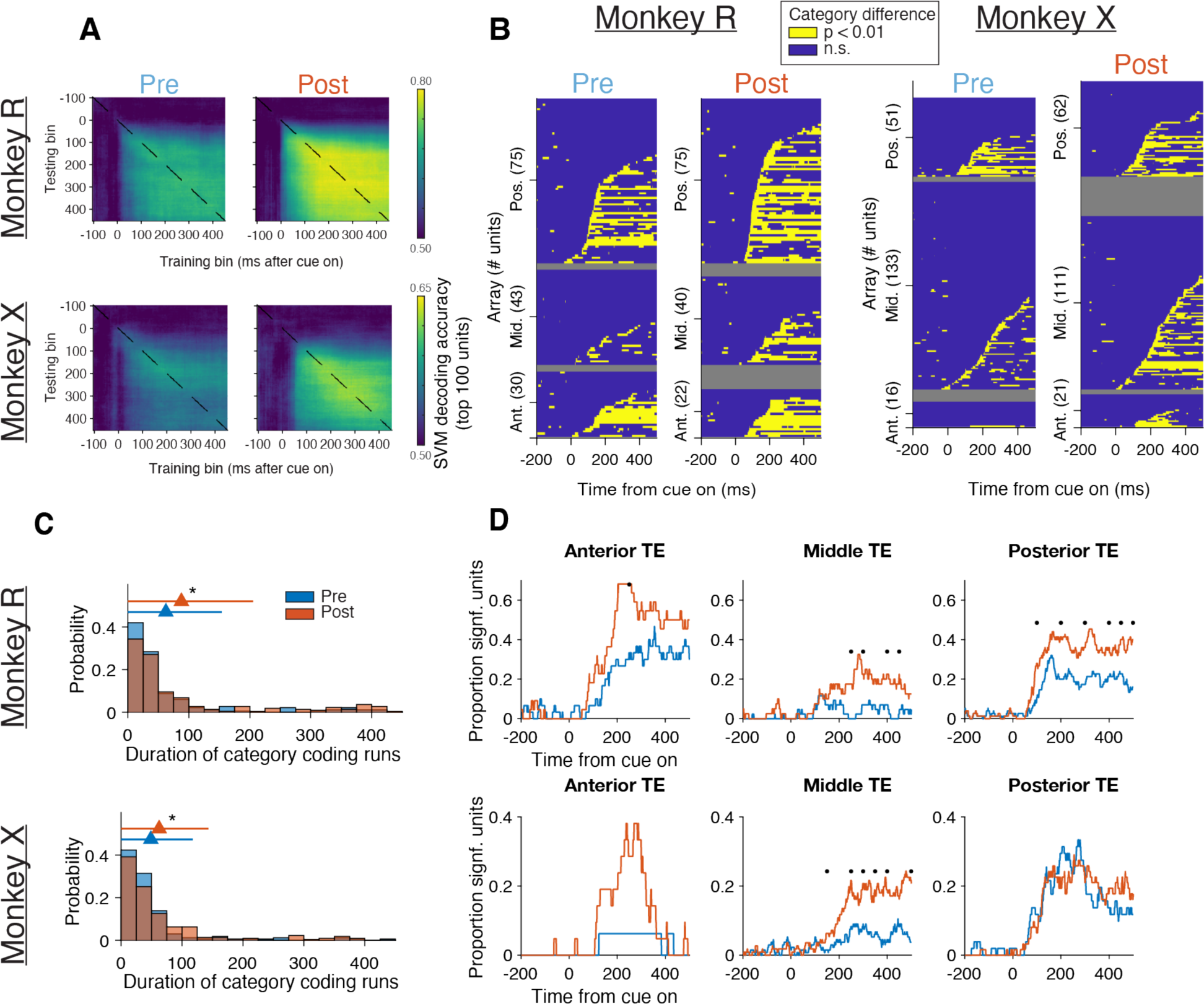
**A,** accuracy of abstract category SVM decoders, trained and tested on neural population response vectors from different timepoints. The same set of top 100 units was used for all train/test combinations. **B,** time-courses of significant difference of category-averaged responses for all units. Each row represents a single unit. Yellow represents significance (see Methods, p < 0.01). **C,** net proportion of units showing a significant difference at each timepoint in **A**. Marks above the data represent significant pre-post difference (two-sided chi-square test, p < 0.05). **D,** probability distribution for the durations of all single unit time courses (not including zero duration) of significant single-unit category coding from the analysis in **B** (p < 0.05, one-sided rank-sum test).

## Discussion

In this study, category training with natural images enhanced neural category coding in area TE. Three factors might have supported enhanced category coding. First, a higher proportion of units coded for category after training (Fig 2D, 3A, 4C), suggesting that new units were likely recruited that hadn’t previously demonstrated category selectivity. Second, single units broadened their responsiveness (Fig 3B,C), and showed a concomitant decrease in sparseness (Supp Fig 4D), responding to larger numbers of cats or dogs in a manner that increased population-level category information. Third, some units increased the duration of their category-selective responses (Figs 4B,D). In line with previous studies, no units strongly responded to a majority of images within a category, with most units responding nearly equally to cats and dogs^8^.

Since the most informative category coding emerged 150-200 ms into TE neurons’ visually evoked responses, and given that the latency of visual activity in TE is around 75 ms, lateral inhibition, feedback processing, or both, are candidate mechanisms for this category-level selectivity. In sum, the improved category coding in TE after learning correlated with improved behavioural accuracy, and was likely driven by a larger fraction of category-responsive neurons, a broadening of single-unit response profiles, and increased duration of category-coding responses.

Did category learning, in and of itself, cause the observed changes in area TE? The evidence suggests that it did: 1) whereas previous studies have relied on extensive periods of training to study neural representations of category or value in TE, we have shown similar effects after approximately one week of training, after which the monkeys were not expert at this task (as evidenced by the reduction in accuracy and increase in reaction times from training to transfer test), arguing against an effect of overtraining or familiarity. 2) The effects of pre-exposure to the stimuli in the baseline period were generally null or smaller than the effects of the category training, again arguing against an effect of familiarity. 3) Since the monkeys were not performing the categorization task when these data were recorded, it must have been a learned association, rather than immediate task salience, that drove these differences in neural responses to the categories. 4) Due to the asymmetric reward structure of our training task, we can’t rule out a role for reward associations in these changes — i.e., the monkeys may have learned to associate dogs with the liquid reward, and cats with the different “reward” of avoiding a delay in the task (see Methods). We observed no non-specific decrease in the sparsity of neural response towards dogs (Supp Fig 4D), and only a mild bias towards dogs in one monkey for overall image selectivity (Fig 3B), arguing against a non-specific reward signal for dogs as the mechanism underlying the observed increases in category information. Further, since it is unlikely that the monkeys would have memorized the 480 test image associations, the monkeys would still have had to visually distinguish cats from dogs via a process of generalization to expect a reward. In conclusion, we propose that visual category learning is, at least partially, supported by enhanced neural representations of category in area TE.

## Acknowledgements

We are grateful to Drs Fernando Ramirez, John H Wittig, Bing Li, Wenliang Wang, and Reona Yamaguchi for helpful feedback while drafting this manuscript.

## Methods

### Experimental subjects / housing / care

Experiments were performed with two Japanese monkeys (*Macaca fuscata*) that were provided by the NBRP-Nihonzaru, which is part of the National Bio-Resource Project of the Ministry of Education, Culture, Sports, Science and Technology (MEXT, Japan). Monkey R was a 12-year-old male weighing 9 kg, and Monkey X was a 13-year-old male weighing 11kg. Monkeys were housed in adjoining individual primate cages that allowed social interaction. The monkeys had access to food daily and earned their liquid during and additionally after neural recording experiments on testing days. Monkeys were tested 5 days per week. All surgical and experimental procedures were approved by the Animal Care and Use Committee of the National Institute of Advanced Industrial Science and Technology (Japan) and were implemented in accordance with the “Guide for the Care and Use of Laboratory Animals” (eighth ed., National Research Council of the National Academies).

### Surgery

Each monkey was first implanted with a titanium head holder approximately four (Monkey R) or two (Monkey X) months prior to electrode implantations. Three microelectrode arrays (Utah arrays, iridium oxide, 96 electrodes, 10 *×* 10 layout, 400 *µ*m pitch, 1.5 mm depth, Blackrock Microsystems, Salt Lake City, USA) were surgically implanted in the anterior, middle, and posterior parts of area TE in the left hemisphere for Monkey R, and four arrays, one additionally in area TEO (data not shown), were implanted in the left hemisphere for Monkey X. Surgical procedures were similar to those described previously^10^, and the procedures for Monkey X have been previously published^11^. For both monkeys, a bone flap located over the temporal cortex was temporarily removed from the skull and a CILUX chamber was placed onto the anterior part of the skull to protect connectors of the arrays.

#### Behavior

##### Behavioral testing

All behavioral tests were carried out using a shielded room. Monkeys were seated in a primate chair, and responded with a touch-sensitive bar that was mounted on the chair at the level of the monkey’s hands. The display was a 21-inch color CRT monitor (GDM-F520, SONY, Japan), and the center of the monitor was placed 56.6 cm in front of the monkey’s eyes. The total reward delivered in each session was about 120 ml of juice for Monkey R and about 420 ml of water for Monkey X. The monkeys were head-fixed for all behavioral testing, and eye-tracking was performed with an infrared pupil-position monitoring system (iRecHS2, Matsuda; http://staff.aist.go.jp/k.matsuda/iRecHS2/). The visual stimuli were presented using the Matlab (Mathworks) Psychtoolbox (Kleiner et al., 2007) on Windows operating system (Microsoft). Task control was performed by the REX real-time data-acquisition program adapted to QNX operating system (Hays et al., 1982).

##### Pre-training

The monkeys were first trained to use the touch bar to receive a reward. Then, a red/green color discrimination task was introduced (Bowman et al., 1996). Each trial began with a bar touch, and 100 ms later a small red target square (0.5 x 0.5 degrees visual angle) was presented at the center of the display. Monkeys were required to continue touching the bar until the color of the target square changed from red to green. Color changes occurred randomly 500 - 1500 ms after bar touch. Rewards were delivered if the bar was released between 200 and 1000 ms after the color change; releases occurring either before or after this epoch were counted as errors.

##### Category pre-training

The category pre-training paradigm was similar to the basic red/green task described above, except that now the initial “no-go” period became the second option in a cued two-interval forced choice task. Each trial began when the monkey pressed the bar, and one of two Walsh patterns was presented. After 350 - 400 ms, the red fixation point appeared, and after 1-3 seconds, it turned green. Release in red ended the trial, whereas release in green led to the trial’s associated outcome. The trial outcome was either a timeout (4-6 seconds) or a reward, depending on which Walsh pattern was shown. This outcome pattern led monkeys to release in red for one pattern, to avoid the associated timeout, and to release in green for the other, to collect the reward. (We have previously shown that avoiding delays to reward has subjectively similar value to monkeys as reward itself^12,13^.) Thus, on each trial, the monkeys had to choose between releasing in the red or green period — a two-interval forced choice task. All trials were followed by an inter-trial interval of 1 - 1.1 seconds.

##### Cat-dog training and transfer test

The experimental paradigm and timings for the cat-dog training task were similar to that described for the category pre-training task, except that the two Walsh patterns were replaced with images of cats and dogs. Further, monkeys were required to fixate on a central fixation point to begin the trial, at which point an image appeared. 350 - 400 ms later, the red cue appeared, and monkeys were allowed to break fixation (see Supp Fig 1B). Images of cats were associated with the timeout, whereas images of dogs were associated with rewards. Monkeys therefore learned to release in red for cats and release in green for dogs.

Monkeys were tested on this task with the small, training image set (20 cats and 20 dogs; Supp Fig 1B) until they reached a performance target (80% correct). Then a transfer test was performed with a larger, held-out image set (240 cats and 240 dogs) to assess their ability to generalize. We note that the 520 total images used during cat-dog training are the same 520 images used in all the passive viewing experiments. Thus, the 480 images in the transfer test were not strictly novel, but had never before been categorized by the monkeys.

#### Passive Viewing Task

Monkeys were trained to keep their gaze in the center of the CRT monitor while images appeared at a moderate speed. Monkeys viewed five images on each trial and received a reward at the end (Supp Fig 1A). Each trial began with a fixation point (0.4 degrees of visual angle) that served as an invitation for the monkey to look at the center of the screen. When the monkey’s gaze entered the fixation window (Monkey R: 4 x 4 degrees; Monkey X: 6 x 6 degrees), five images (12 x 12 degrees) were presented serially, each appearing for 350-400 ms, with 350-400 ms of blank screen between images. The fixation point remained present during the entire trial. If the monkey’s gaze exited the fixation window during the trial, the trial was immediately terminated, and the trial with the identical image sequence was repeated. After all five images, a liquid reward was delivered, the monkey was allowed to break fixation, and there was an inter-trial interval of 1 second. 520 images were used in total, with 260 cats and 260 dogs. Images were presented in randomized blocks such that each image was presented once in the block.

We recorded six days of baseline cat/dog passive viewing data for Monkey R and two days for Monkey X. After that, we performed separate, unrelated passive viewing experiments for 2-3 months, during which the neural activity recorded by the arrays qualitatively changed. Thus, we recorded one additional pre-training cat/dog passive viewing session immediately before cat/dog training for both monkeys. After training, we recorded two days of passive viewing for Monkey R (because the first day had too low of a trial count; only 3-4 presentations of each cue, compared with 5-8 for good days – only data from the second day was included in our analyses) and one day for Monkey X (20 presentations per cue).

#### Neural Recording

##### Recording and Spike Sorting

Neural data, task events, and eye positions were recorded using Cerebus^TM^ system (Blackrock Microsystems). Extracellular signal was band-pass filtered (250–7.5 k Hz) and digitized (30 kHz). Units were sorted online before each recording session for the extracellular signal of each electrode using a threshold and time-amplitude windows. The spike times of the units were stored using Cerebus Central Suite (Blackrock Microsystems). Single units were refined offline by hand using principal-component-analysis projection of the spike waveforms in Offline sorter^TM^ (Plexon Inc., Dallas, USA).

##### Data exclusion

Inspection of the data using standard raster plots revealed Day 4 of Monkey R’s baseline data to be contaminated by noise, so it was excluded from the analyses. Similarly, our first attempt at recording a post-training passive viewing session for Monkey R only had 3-4 presentations per image due to the monkey not being as motivated as usual, so we performed a second session the following day, which was used for analysis.

For all analyses of neural data during passive viewing, we used data exclusively from completed trials, that is, trials without fixation errors.

Neurons with less than 20 spikes in a given session were removed from the analysis. Otherwise, all units recorded from TE were used in a given analysis, except where noted (e.g. Figure 2).

#### Data Analysis

##### Behavioral Analyses

To compute the fraction of trials correct by session sextile, sessions were split into six equal blocks of trials, including incomplete trials (i.e. including fixation errors and pre-cue bar releases), and the fraction correct trials was computed within each block. Reaction times for correct cat trials (Fig 1C, iv) were calculated as the time between when the red cue appeared and when the bar was released.

##### Image Analysis

To extract the foreground of images, we took advantage of the fact that they were cropped onto white backgrounds. We k-means clustered the RGB values into 4 clusters, which consistently reported the background as one of the colors. We subsequently set the pixels belonging to any other cluster to black, and then performed an image fill operation to fill holes. The number of foreground pixels was the number of black pixels after this operation. HSV distributions were computed from these foreground pixels. The image-mean luminance was calculated using a standard RGB conversion^14^. Spatial frequency energy was calculated using a 2-D Fourier transform of the image and summing the energies across all orientations at each frequency.

##### Initial Neural Analysis

To select the most informative analysis window, we took an unbiased approach using Monkey R’s data (which was acquired first) and an SVM classifier with an “abstract category” approach (see below) (Supp Fig 2A)^6^. The classifier’s accuracy was assessed at 25 ms intervals, with window starting points ranging from 50 ms before image onset to 300 ms after image onset. Window sizes varied from 50 ms to 300 ms in 25 ms increments. Longer windows were more informative without exception. The accuracy of the decoders using the smallest window peaked at 150 ms. As the latency of visual information arriving in TE is roughly 75 ms. we chose 75-175 ms and 175-275 ms as two spike windows to correspond roughly to the first wave of activity in TE, and a non-overlapping window with better decoding. In analyses of Monkey X’s data, similar trends with respect to these analysis windows were observed, so we kept the windows the same.

##### Decoding Analysis

For the SVM analyses (Figure 2B,C,D), we asked how well the neural population response (spike counts) could decode the category (cat or dog) of the presented image. N neurons’ spike counts (predictors) provided a response vector for each of the P image presentations (observations), so the input matrix to the SVMs had dimension P x N. In order to more accurately assess category information in the population, we used “abstract category decoding”^6^, which prevents overfitting to particular images by using different images for training and test data. Images were divided into five subsets, and each set of image presentations corresponding to each subset was used once as a held-out set. Using the MATLAB function ‘fitclinear’ with lasso regularization, we discovered that the “sparsa” solver, which only uses up to 100 features, delivered better abstract category decoding than the “sgd” solver, which uses an arbitrary number of features. Thus, in order to maximize our estimates of decoding accuracy, we limited our decoding to using the top 100 category-coding single units. To rank the single units by individual category coding, we ran 1-D SVM decoders on each unit (i.e., P x 1 inputs to the decoder), and ranked units based on the resulting cross-validated accuracies. Population decoding was repeated using different 100 ms wide time bins (Figure 2B); or across sessions (Figure 2C); or adding the top 100 units one at a time (Figure 2D).

To fit the sigmoidal curves in Figure 2D, we used the MATLAB function ‘fittypè with the sigmoidal equation ‘a1/(1 + exp(-b1*(x-c1))) + d’. The first three points were excluded to achieve a stable fit for the sigmoidal curve (i.e. positive amplitudes, half-max x-values inside the domain of the data).

For the time-swap analysis (Figure 4A), decoders were trained as above, but then evaluated on spike-count data from all other bins.

##### Single-unit Analyses

For the GLM analysis (Figure 3A), for each neuron, we asked if that neuron fired at significantly different rates for cats and dogs. Spike counts for each image presentation were regressed against image category with a simple linear model, log(spike count) ∼ image category (Poisson link function). Units were counted as having a cat/dog difference if they had a significant coefficient in the model (p < 0.05).

For the image responsiveness analysis (Figure 3B), for each neuron, we asked if that neuron fired above its baseline rate for each image. A one-sided t-test was used to compare spike counts in the window 175 to 275 ms after image onset to those in a pre-onset window of the same size (-150 to -50 ms), and t-tests with p < 0.05 were considered significant. The fraction of images of each category to which each neuron significantly responded was plotted. Each neuron’s category selectivity was then quantified as the absolute difference between the fraction of cat and dog images to which it responded (Figure 3C).

To fit the slope of the population’s responsiveness (Figure 3B), a line was fit using major-axis regression^15^. Major-axis regression is suitable when there is no independent/dependent variable pairing, as is the case here; the two variables are equivalent and we do not *a priori* expect a causal link in one direction or the other. Major-axis regression was performed analytically (i.e. with an explicit formula, using the MATLAB function maregress), returning the line’s slope and the corresponding 95% confidence interval (Figure 3D).

For the time-course analysis of category differences (Figure 4), smoothed, normalized category-level firing rates were first estimated for each neuron using a kernel density estimate (Gaussian kernel, bandwidth 20 ms) applied to the collated peri-stimulus spike times from all the cat or all the dog image presentations. The densities (units of 1/sec) were then converted to average rates by multiplying them by the total number of considered spikes and dividing them by the number of considered trials (units of spikes/trial/sec). The two spike rates were subtracted and the absolute value was taken as the category difference over time. To assess statistical significance of this difference, the same analysis was repeated 50 times with shuffled data, and timepoints where the real difference (at that x-value) exceeded every shuffled data point (at that x-value) were considered to have significant category difference at that timepoint (Figure 4B). We then calculated the fraction of units with significant category difference at each timepoint and compared pre vs post-training sessions (Figure 4C). Significance was assessed with a chi-squared test at 50 ms intervals. Finally, the durations of periods of continuous significant category difference were calculated for each session, and the distribution of run lengths pre vs. post were compared with a one-sided ranksum test.

To quantify the sparseness of neural responses, two previously described non-parametric metrics were used^8^. Briefly, the “sparseness index” quantifies the extent to which a neuron’s total firing is distributed amongst few or many images in a set.

---

**Supp Fig 1:**
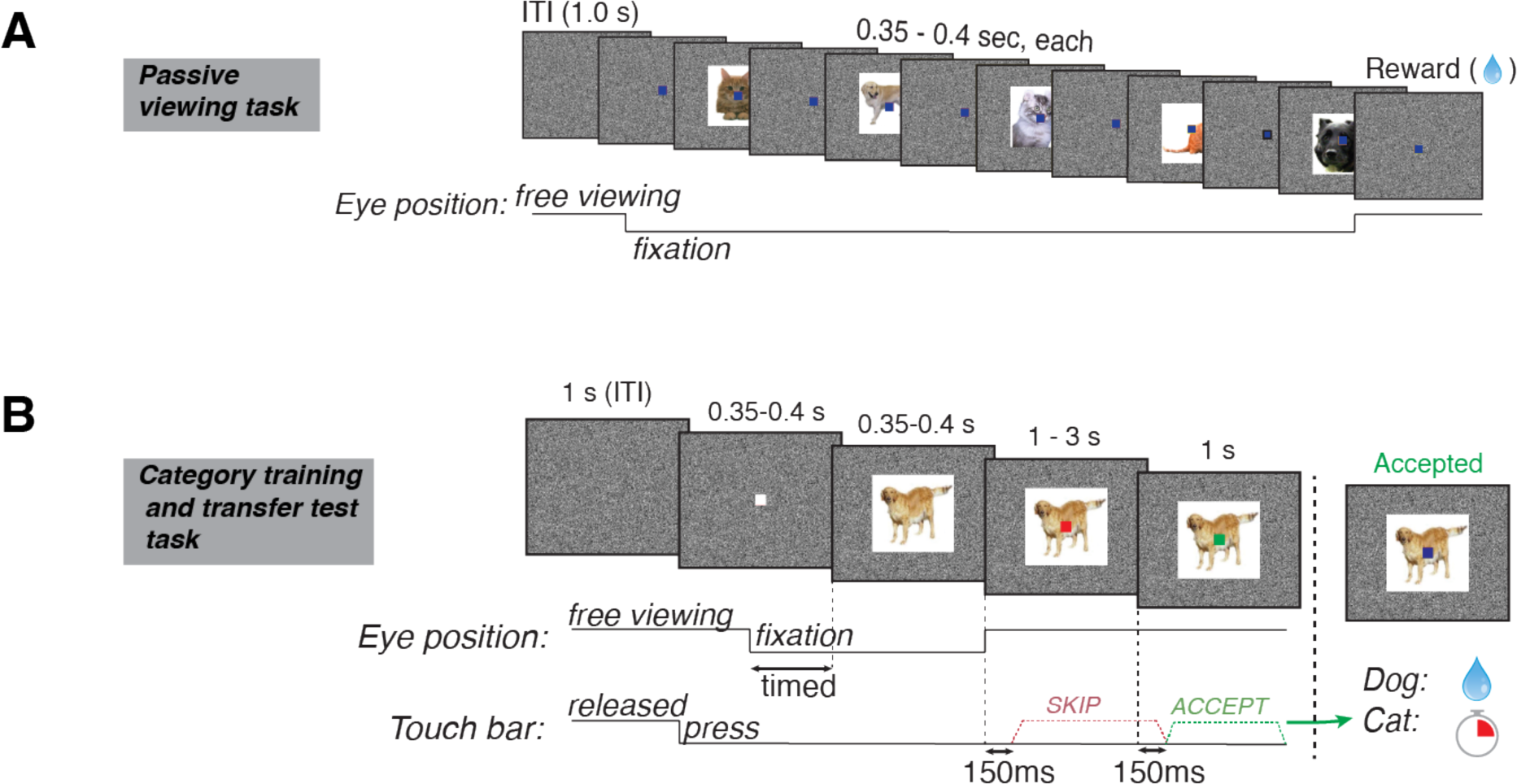
**A,** diagram of the passive viewing task. In brief, monkeys were required to fixate on the central fixation square while five images were presented in succession, followed by a liquid reward and an inter-trial interval. See methods for details. **B,** diagram of the category training and transfer test task. In brief, in each trial, monkeys were required to release a touch-bar in one of two intervals to indicate whether the image on screen was a cat or a dog. To correctly respond to a cat stimulus, monkeys had to release during the interval in which the central fixation square was red, which allowed them to avoid a timeout; to correctly respond to a dog stimulus, monkeys had to hold the bar through the red period and release when the square turned green, which resulted in a liquid reward. See Methods for details.

**Supp Fig 2:**
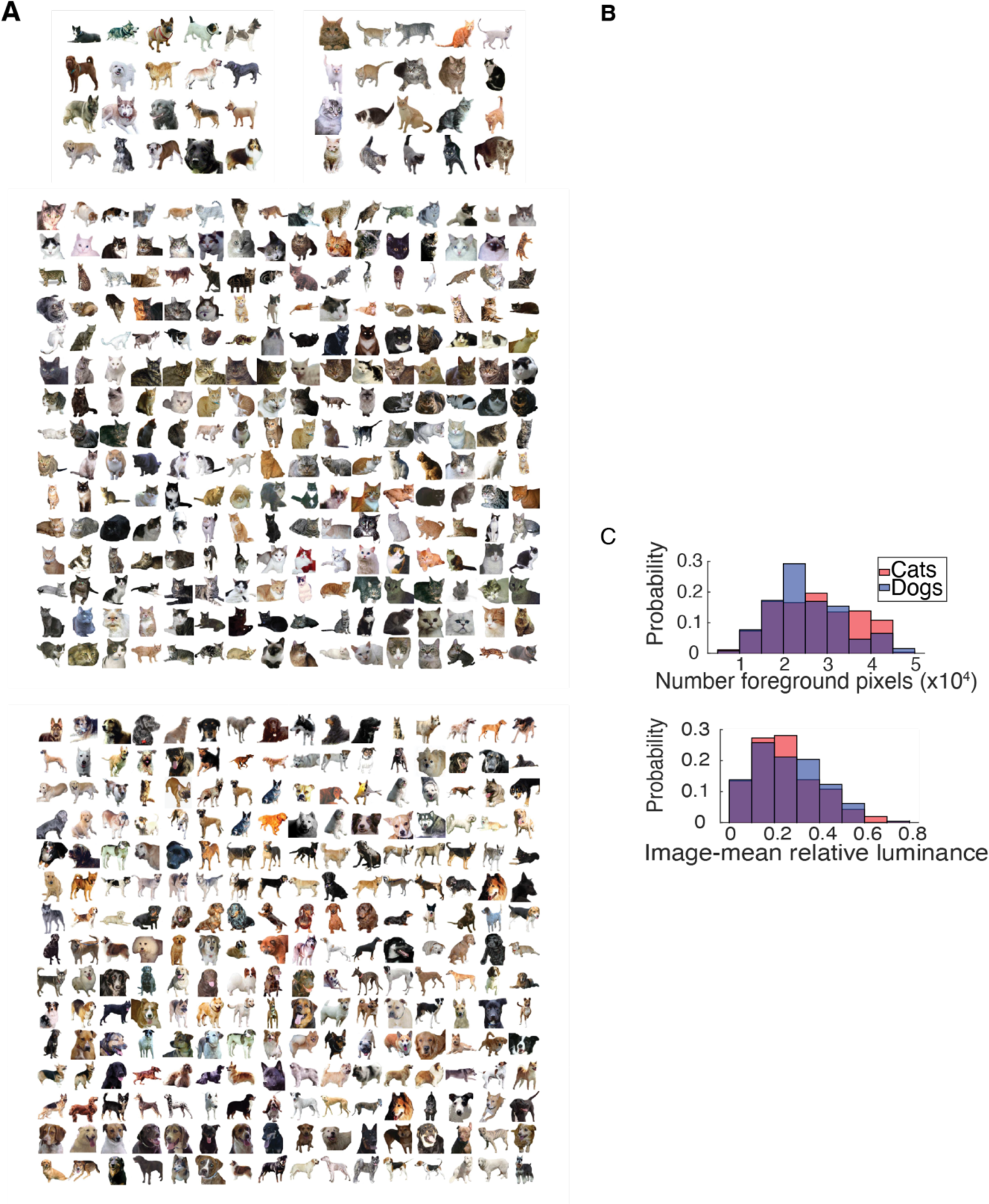
**A,** the 520 stimuli used in this experiment, broken down into the 20/20 and 240/240 sets used for behavioral training and testing, respectively. Images were natural images with the background removed. **B,** distribution of the colors of pixels in the cat and dog datasets. Only foreground pixels were considered. The foreground was extracted using a k-means clustering algorithm (k=6) and removing the white component. **C,** analysis of object size (number of foreground pixels), image-mean relative luminance, and frequency components of the cat and dog stimuli.

**Supp Fig 3:**
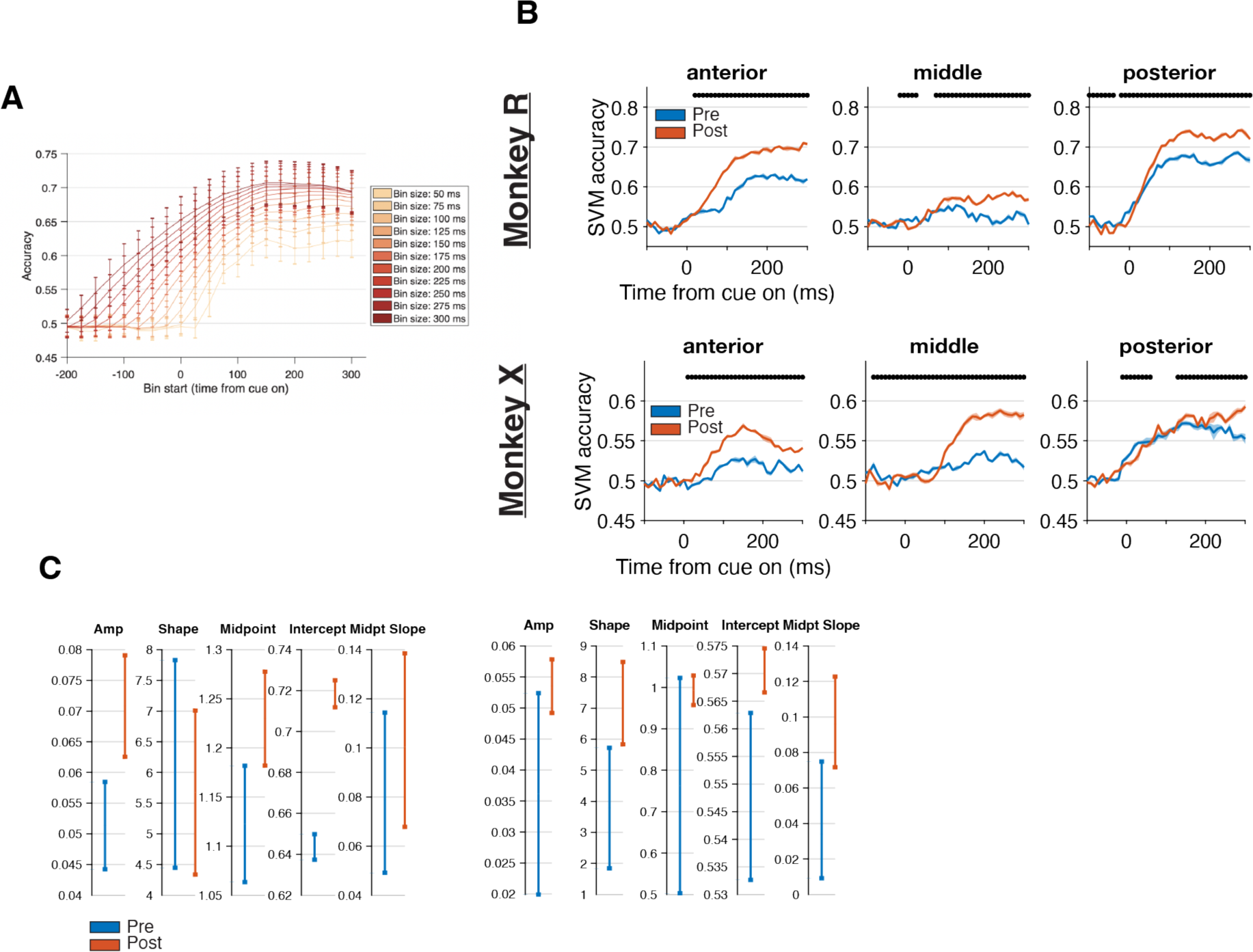
**A,** time-course analysis using abstract category SVM decoders, with spike counts collected over different bin widths. **B,** time-course of abstract category SVM decoders (as in Fig 2b), but using the top 100 units (or all units, whichever was fewer) from each separate Utah array. **C,** parameter means and 95% confidence intervals for sigmoidal fits shown in Figure 2D.

**Supp Fig 4:**
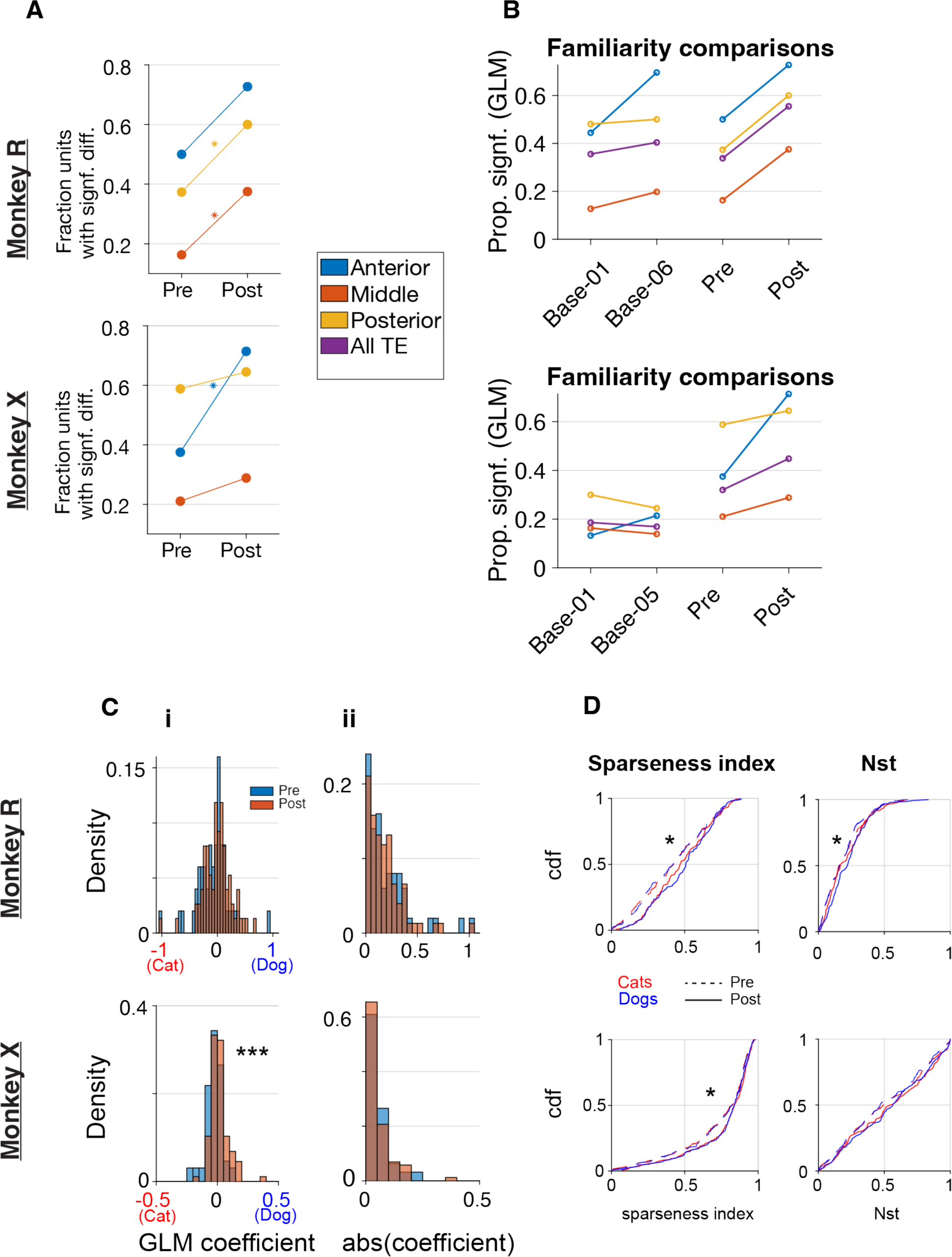
**A,** fraction of single units significant in a GLM regressing image category vs. spike count 175-275 ms after image onset (as in Fig 3a), broken out across the Utah arrays. **B,** the same analysis as in A, but comparing the first and last baseline days to the pre- and post-training days. **C,** distributions of GLM coefficients broken out by pre vs. post-training days. i, the signed value of the coefficients, showing a slight bias towards dog-preferring units in Monkey X (***, p < 0.001, two-sided ranksum test); ii, the same data as in i, plotted with its absolute value to demonstrate no increase in pre vs. post-training absolute spike rates (n.s., one-sided ranksum test, p > 0.05). **D,** cumulative distributions (across all single units) of sparseness index and Nst^8^, broken out by cat vs. dog and pre vs. post-training days, showing a decrease in sparseness (increasing sparseness index and Nst; *, one-sided ranksum test, p <= 0.05).

**Supp Fig 5:**
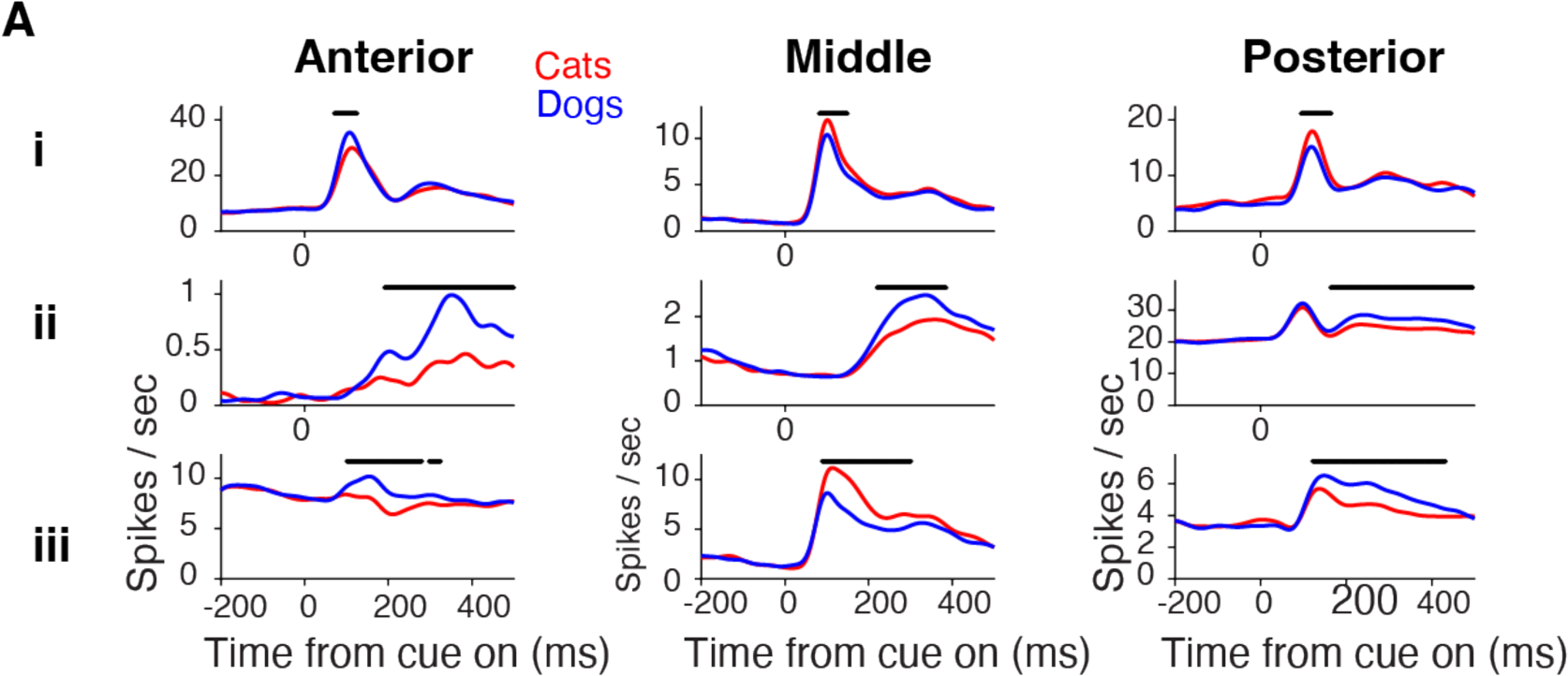
**A,** smoothed mean responses to cats and dogs for representative units across arrays. Black lines above the data indicate significant cat-dog difference (p < 0.01 via bootstrapping) (as in Fig 4a). Units had diverse response patterns, showing significant category differences during either **i)** only the initial wave of visually-evoked activity; **ii** ) the sustained portion of visually-evoked activity; or **iii**) both.

